# A novel reverse lipase toxin substrate of the *Staphylococcus aureus* type VII secretion system

**DOI:** 10.64898/2026.06.22.733114

**Authors:** Andrew B. Higginson, Jasmine Soh, Stephen R. Garrett, Terry K. Smith, Tim R. Blower, Tracy Palmer

## Abstract

The type VII secretion system (T7SS) is found in many Gram-positive bacteria and secretes toxins with antibacterial activity. Most characterised substrates have an N-terminal LXG domain that interacts with other helical partner proteins to form a composite T7SS targeting signal. Here we describe only the second substrate family to have a reverse domain arrangement. We show that TslM has a C-terminal LXG-like domain and an N-terminal lipase domain that has phospholipase activity. Secretion of TslM requires a single helical partner protein that binds to the TslM C-terminus, and its toxic activity is neutralised by a distinct family of membrane proteins. Genome analysis reveals that *Staphylococcus aureus* strains have the capacity to encode up to seven paralogous copies of this toxin family. Taken together our findings show that lipases are an important component of the staphylococcal T7SS toxin arsenal, and that toxins with a reverse domain arrangement are more widespread than previously appreciated.

## Introduction

*Staphylococcus aureus* is a Gram-positive organism that is closely associated with humans and other mammals. It is persistently carried by up to 30% of the human population, with the nares as the primary colonisation site. *S. aureus* carriage is asymptomatic, and the bacterium is considered part of the nasal commensal flora. However, it can also cause severe disease including bacteraemia and endocarditis if the skin or mucosal barrier is breached.

The composition and stability of microbiomes is influenced by co-operative and competitive interactions between its members. Numerous mechanisms have been identified that underpin interspecies interactions. Co-operation is frequently facilitated through shared resources such as metabolites and essential cofactors (Bilici et al., 2025; Blasche et al., 2021; Kramer et al., 2020). Competitive interactions may be mediated by bioactive small molecules such as antibiotics, ribosomally synthesised peptide-based bacteriocins, or larger proteinaceous toxins (Clardy et al., 2009; Cotter et al., 2012; Heilbronner et al., 2021). Within the nasal microbiome, *S. aureus* is antagonised by lugdunin, a non-ribosomally synthesised peptide antibiotic produced by *Staphylococcus lugdunensis* (Krismer et al., 2017; Zipperer et al., 2016) and epifadin, an antimicrobial peptide polyene produced by strains of *Staphylococcus epidermidis* (Torres Salazar et al., 2023). The production of these molecules may explain the negative association between *S. aureus* presence and either of these two Staphylococcal species in human nasal microbiomes (Rosenstein et al., 2024; Torres Salazar et al., 2023).

*S. aureus* strains are equipped with numerous mechanisms that allow them to compete with other microorganisms. A handful of strains encode bacteriocins, usually on plasmids, which are active against closely related Gram-positive bacteria (Garrett & Palmer, 2024). Some strains also produce a novel antibacterial protein (termed ABP) that requires processing by a cognate secreted protease for full toxic activity (Colautti et al., 2025). In addition to these sporadically-encoded competitive factors, all *S. aureus* strains possess the type VII secretion system (T7SS) alongside variable numbers of genes coding for T7-secreted toxins (Bowman & Palmer, 2021; Warne et al., 2016).

The T7SS is widely distributed among Gram-positive bacteria and Actinomycetota (Garrett et al., 2024). The core components of the T7SS are a membrane-embedded ATPase TsxC (termed EssC in *S. aureus*), at least one secreted helical hairpin protein of the WXG100 family (TsxA; EsxA in *S. aureus*) (Shah et al., 2026), and TsxB (*S. aureus* EsaB), a small protein with a ubiquitin-like fold. Alongside these conserved components, additional suites of system-specific membrane proteins are associated with the T7SS across the different bacterial phyla, leading to sub-classification of the T7SS from T7SSa through to T7SSj (Garrett et al., 2024). *S. aureus* harbours the T7SSb variant, which it shares with some other Bacillota members including *Streptococcus* (Whitney et al., 2017), *Listeria* (Bowran et al., 2023; Bowran & Palmer, 2021) and *Bacillus subtilis* (Baptista et al., 2013; Huppert et al., 2014). Additional membrane components of the T7SSb are EsaA, EssA and EssB (Fig 1A, 1B).

**Figure 1.**
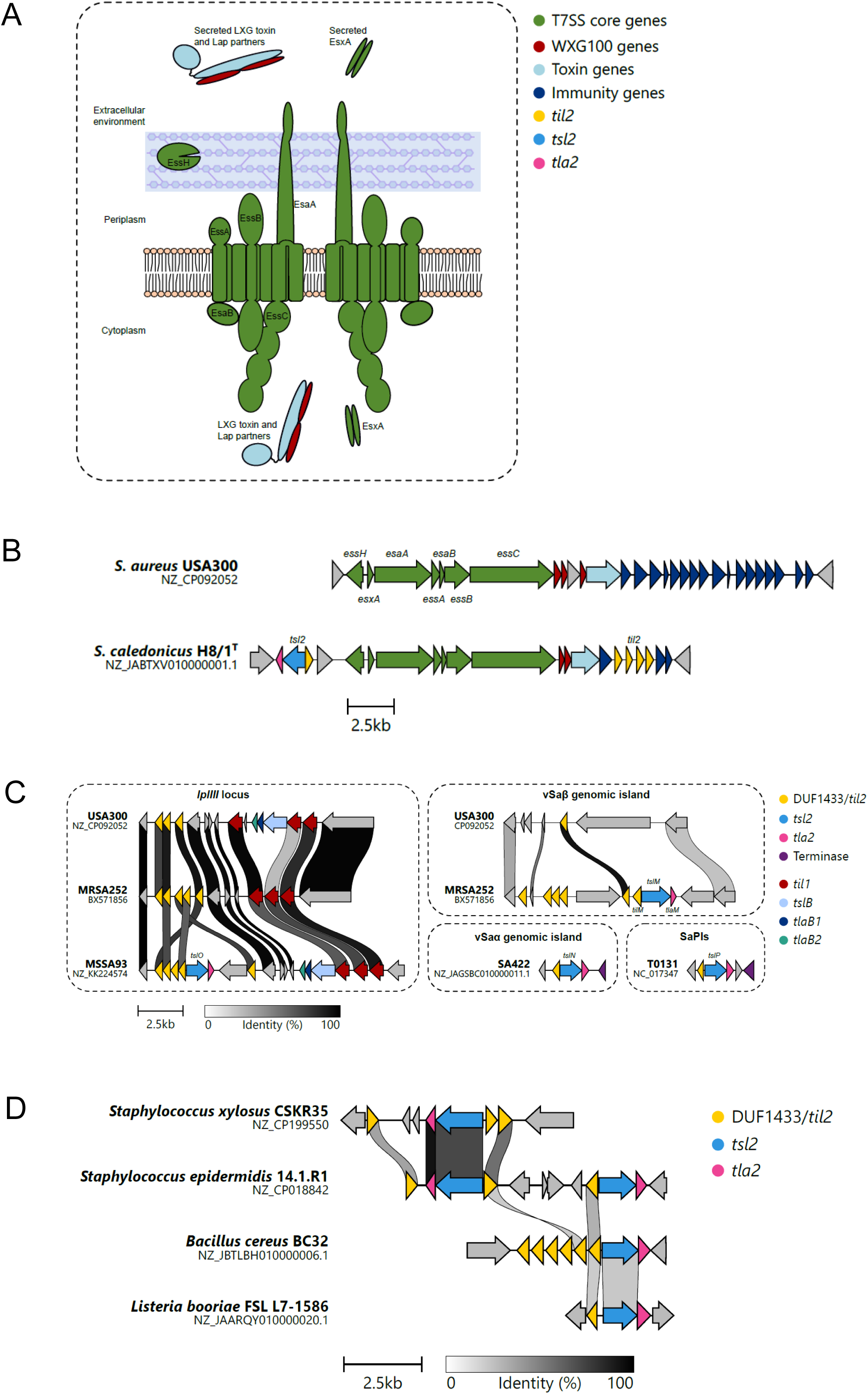
The Staphylococcal T7SS and the Tsl2 Family of T7SS lipase substrates. A. Cartoon representation of the T7SSb found in *S. aureus* and other staphylococcal strains. Core components of the secretion system are shown in green, toxin substrate in blue and Lap secretion partners in maroon. EssH is a peptidoglycan hydrolase that is required for assembly of the T7SS (Agyen et al., 2026; Bobrovskyy et al., 2018). B. The T7SS-encoding loci in *S. aureus* USA300 and *S. caledonicus* H8/1^T^. C. Tsl2 proteins can be encoded in four *S. aureus* genomic contexts; (*left*) the *lplIII* locus, (*right*) νSaβ genomic island νSaα genomic island, and SaPIs, of which one is shown. Variable numbers of *til2* genes (yellow) are present at the *lplIII* locus and νSaβ island of USA300 and MRSA252. The shaded links indicate genes with high similarity. Genes shown in grey are unrelated to Tsl2. D. Tsl2 proteins can be found encoded in other Bacillota, including the *Staphylococcus*, *Bacillus*, and *Listeria* genera. Example loci encoding Tsl2 homologues (WP_107544516.1, WP_102841626.1, WP_098326801.1 and WP_187125958.1, respectively) are shown in blue, with shaded links indicating genes with high similarity. Note the presence of two different Tsl2 family homologues encoded in *S. epidermidis* 14.1.R1.

To date, the best characterised substrates of the T7SSb are antibacterial toxins that harbour an LXG domain at their N-terminus and a toxin domain at the C-terminus (Alcock et al., 2025; Ulhuq et al., 2020; Whitney et al., 2017). Structural studies have shown that the LXG domain adopts a helical fold and that it binds two or three small helical partner proteins, termed Laps (for LXG-associated α-helical proteins) to form an extended α-helical rod (Alcock et al., 2025; Klein et al., 2024; Yang et al., 2023). Targeting sequences present in the helical rod, including the LXG motif of the toxin and an FxxxD motif or variant at the C-terminus of one of the Lap partners are critical for secretion of the toxin-Lap complex (Anderson et al., 2013; Klein et al., 2022, 2024). The C-terminal toxin domains are highly variable, with activities such as DNase, lipid II phosphatase and NADase (Cao et al., 2016; Whitney et al., 2017). LXG toxins have been shown to mediate inter- and/or intraspecies competition, and strains encoding these toxins also produce cognate immunity proteins for self-protection (Cao et al., 2016; Job et al., 2024; Kobayashi, 2021; Ulhuq et al., 2020; Whitney et al., 2017).

Recently, a novel family of T7SS substrates, the Tsl1 family, was described in *S. aureus* and other Bacillota (Garrett et al., 2023). These proteins show an unexpected reverse domain organisation compared to other known T7SS substrates, having a toxic lipase domain at the N-terminus and an LXG-like domain at the C-terminus. *S. aureus* strains may encode up to three non-identical Tsl1 proteins, TslA, TslB and TslC, at distinct loci. Despite the unusual domain arrangement, TslA was shown to adopt an overall similar architecture to LXG toxins and required interaction of the C-terminal LXG-like domain with two helical Lap-like partner proteins, TlaA1 and TlaA2 for secretion. Tsl1 proteins are co-encoded with clusters of genes encoding Til1 lipoproteins. The canonical Til1 family member, TilA, was shown to bind tightly to the TslA lipase domain, blocking activity and providing protection from self-intoxication (Garrett et al., 2023).

In this study we identify Tsl2, a second family of ‘reverse’ T7SS substrates that can be present in multiple paralogous copies in *S. aureus* strains. Through characterisation of the TslM family member we show that Tsl2 proteins are phospholipase toxins that are dependent on a single Lap partner for secretion and that are neutralised by the DUF1433 family of membrane proteins.

## Results

### Identification of the Tsl2 family of T7SSb substrates

T7SS substrates in *S. aureus* are encoded at conserved loci, usually alongside genes for Lap partners, and for multiple, non-identical copies of immunity proteins. The gene adjacent to the toxin substrate most likely encodes the cognate immunity protein, and it is assumed that at least some of the other copies offer protection *in trans* from polymorphic toxin variants produced by other *S. aureus* strains (Cao et al., 2016; Garrett et al., 2023; Ulhuq et al., 2020; Warne et al., 2016). These loci are hotspots for genetic recombination that can alter the sequences and number of immunity genes (Garrett et al., 2022). Toxin genes may also be absent or frameshifted at these loci in a strain-specific manner, but immunity genes are almost always still retained.

While analysing *S. aureus* strains for the presence of *tslA* and its homologues, we noted strings of genes coding for proteins of the DUF1433 family downstream of *tslB* at the *lplIII* locus in strain USA300 (Fig 1C, left panel). The DUF1433 proteins encoded at this locus are of non-identical sequence, but each is predicted to have an N-terminal transmembrane signal anchor and an extracellular globular domain. We also noted that the number of DUF1433 genes encoded at this locus was variable between strains (Fig 1C, Fig S1). From these features we reasoned that they may be candidates for a novel family of immunity proteins, but were unable to identify a candidate toxin at the locus in either USA300 or MRSA252 strains. To identify a corresponding toxin, we undertook gene neighbourhood analysis using webFlaGs across all available *S. aureus* genome sequences available in RefSeq, with the DUF4133 protein, WP_001034756.1, from USA300 as our input sequence (Fig. S1). From this we found that some strains, including MSSA93, encode additional genes at this locus. Analysis of these revealed an open reading frame specifying a putative phospholipase, as well as a gene encoding a WXG100 family protein (Fig 1C).

The putative phospholipase (WP_033856953.1) has a predicted two domain structure, with the phospholipase domain at the N-terminus, and a predicted helical C-terminal domain. The domain organisation is reminiscent of the ‘reverse’ T7SSb substrate TslA (YP_498992.1), and indeed the phospholipase domains of the two proteins share 22% sequence identity (Fig S2). As TslA homologues can be encoded at three distinct *S. aureus* chromosomal loci, we undertook a BLASTp search using WP_033856953.1 against all *S. aureus* genomes available on RefSeq, identifying a further 316 homologous proteins, ranging in size from 60-546 residues, the majority of which were annotated as predicted lipases. Elimination of obvious truncations below 300 residues resulted in a final list of 305 sequences which clearly represent multiple distinct groups of homologous proteins, based on their differing length and identity to the query. A further webFlaGs analysis revealed that these homologous proteins grouped into four distinct classes based on their sequence and locus, but all shared adjacency to at least one DUF1433-encoding gene and a WXG100 protein-encoding gene (Fig. 1C, Fig S3). The four distinct classes are encoded at the following loci: the νSaβ genomic island, νSaα genomic island, *lplIII* locus, and finally on *S. aureus* pathogenicity islands (SaPIs), which can themselves insert at up to four distinct chromosomal loci. We also noted that a copy of this locus is found adjacent to the T7SS structural genes in *Staphylococcus caledonicus* (Fig 1B). For these and other reasons outlined below, we named this family of predicted novel lipase toxins the Tsl2 family (for <u>T</u>ype VII <u>s</u>ecreted lipase 2) and the lipase genes *tslM,N,O* and *P*, respectively (Fig S4). From this analysis, the majority (64%) of *S. aureus* Tsl2 proteins encoded in RefSeq sequence are TslM, with TslN being least abundant (Table 1).

**Table 1:**
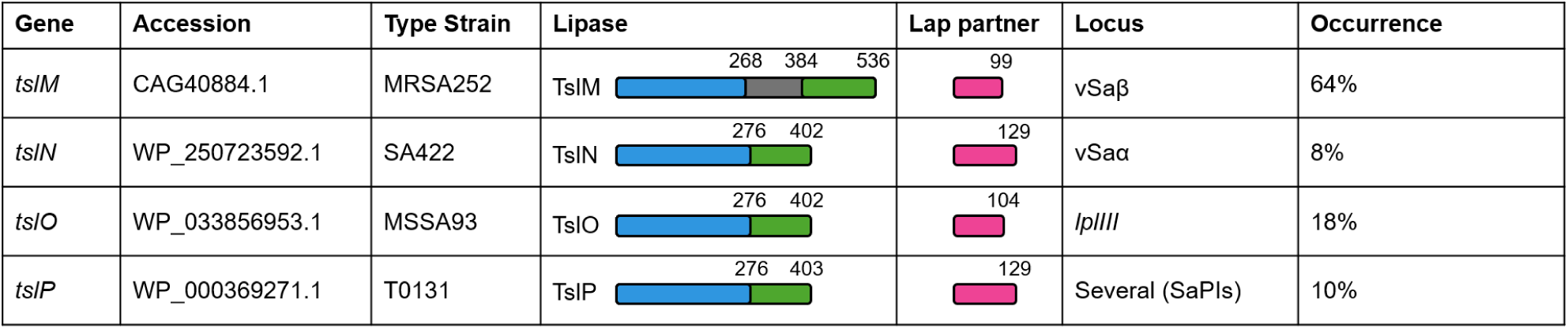
Analysis of *tsl2* family genes found across *S. aureus* genome sequences present in the RefSeq database. Lipase and partner protein domain diagrams are represented to scale, and coloured as follows: lipase domain (blue), LXG domain (green), linker (grey), WXG100 (pink). DUF1433 immunity genes do not differ greatly in length or domain organisation between families, and so are not shown. Occurrence is calculated as a percentage of 305 sequences, based on flanking gene analysis.

We also investigated the prevalence of this novel candidate T7SSb toxin in other Gram-positive species. After adjusting BLASTp searches to exclude *S. aureus*, homologues of TslM were identified in other staphylococci, including *S. caledonicus* (Fig 1B), *S. xylosus* and *S. epidermidis* (Fig 1D). Homologues could also be identified in other genera known to possess the T7SSb, including *Bacillus* and *Listeria* (Fig 1D). These Tsl2 homologues, however, resembled the smaller members of the Tsl2 family (TslP and TslN in *S. aureus*), with an average length of 400aa and no clear predicted middle linker domain, which appears to be unique to the staphylococcal Tsl2 proteins. Smaller Tsl2 protein-encoding genes could also be found in *S. epidermidis*, where strains such as 14.1.R1 can encode both a homologue of TslM and TslP in close proximity (Fig 1D). Interestingly, just as TslP is encoded on SaPIs, this *S. epidermidis* TslP homologue is encoded on SePIs, the equivalent mobile element (Madhusoodanan et al., 2011).

### Characterisation of TslM

As TslM is the most abundant Tsl2 protein encoded in *S. aureus* genomes, we chose to focus on this member of the family. In strain MRSA252, TslM is encoded by *SAR1895*. Six DUF1433 proteins, candidate immunity proteins for TslM, are encoded divergently to *tslM* (Fig 1C). We have named the DUF1433 proteins the Til2 family, with the one coded adjacent to *tslM* as TilM (<u>T</u>ype VII immunity to lipase <u>M</u>) as we show below that this neutralises TslM toxicity. Typical of previously observed immunity rafts, the remaining Til2/DUF1433-encoding genes share varying degrees of similarity to *tilM*, with one being a pseudogene. We named the WXG100 family protein encoded alongside TslM as TlaM (for <u>T</u>ype VII lipase <u>a</u>ssociated <u>M</u>). Also situated on the genomic island are genes encoding a hyaluronate lyase *hysA*, the serine proteases *splC*–*splF* and the *se* enterotoxins (Baba et al., 2008; Moon et al., 2015) (Fig S5). The νSaβ genomic island is classified into twenty-two subtypes (Kläui et al., 2019; Schwendimann et al., 2021), of which *tslM* is encoded on six (III, VI, VIII, IX, XVI and XVII). However, an “orphaned” copy of Til2 is encoded on the majority even when *tslM* and *tlaM* are not present (Fig S5). This island is implicated in both virulence (enterotoxins and leukotoxins) and antibacterial competition, with the *bsa* operon responsible for lantibiotic synthesis (Daly et al., 2010).

TslM is a 536 amino acid protein, with the N-terminal globular predicted lipase domain encompassing aa 1 - 268 and the C-terminal α-helical domain covering aa 384 – 536 – this domain resembles the trafficking domain of other LXG toxins (Fig 2A, Fig 2B, Fig S6). The lipase domain of TslM aligns well structurally with the other characterised lipases including Lip2 (Pignède et al., 2000), with a clearly identifiable serine-aspartate-histidine catalytic triad (Fig 2B inset) and a GXSXG motif surrounding the likely catalytic serine (Fig. S7). Unlike TslA, TslM also has an extensive middle linker (aa 268 - 384) which AlphaFold3 predicts to be unstructured. A similar lack of structure is also observed in the middle region of the T7SS nuclease toxin EsaD (Yang et al., 2023).

**Figure 2.**
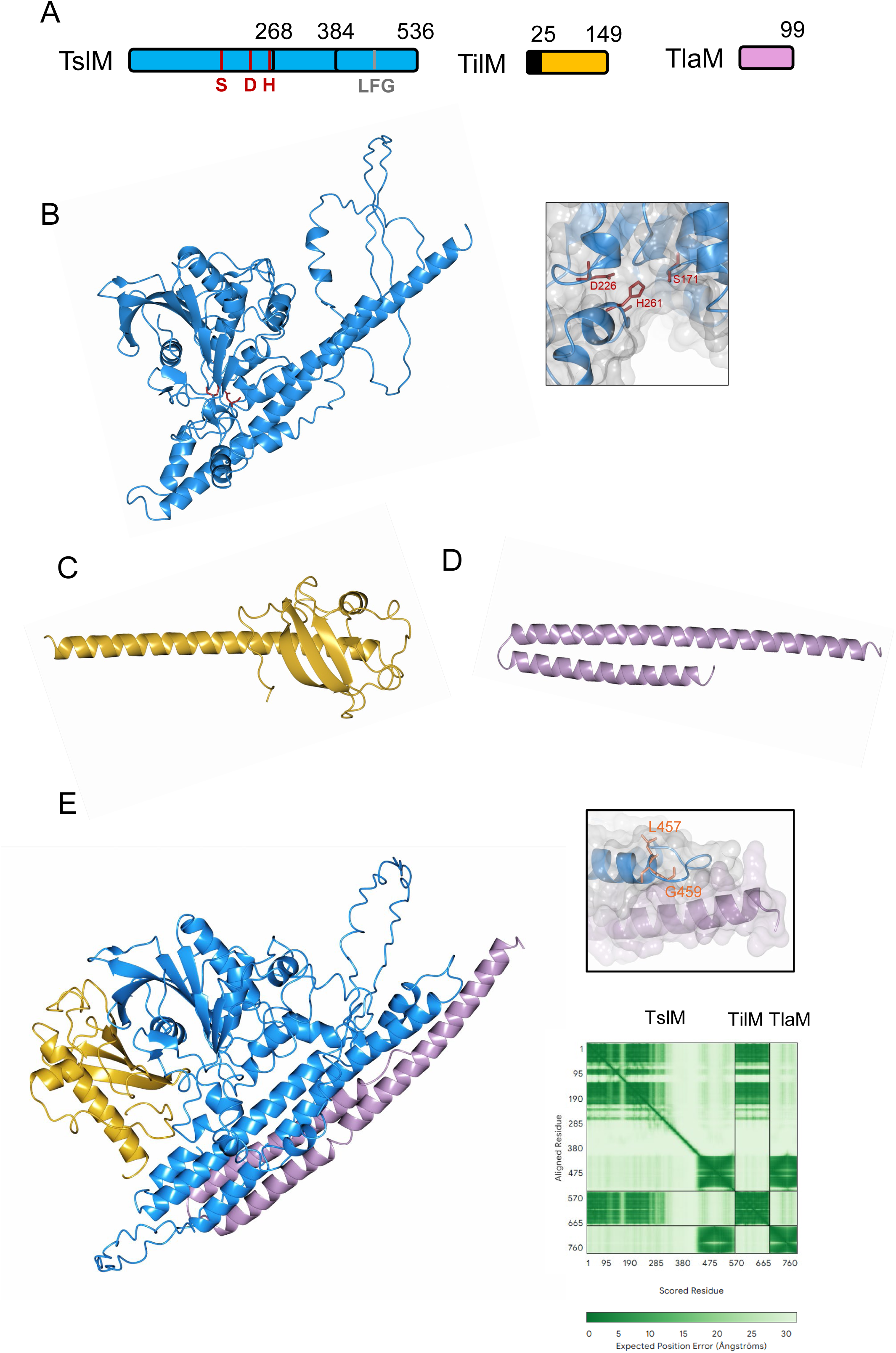
Structural models of TslM, TlaM and TilM. A. To scale domain diagrams of TslM, TlaM and TilM. For TslM, active site residues marked in red and predicted LFG secretion motif marked in grey. For TilM the transmembrane helix is shown in black and the DUF1433 extracellular domain in gold. B. AlphaFold3 model of TslM, pTM = 0.59. Inset: The predicted active site of TslM, with catalytic triad residues shown in red. B. AlphaFold3 model of TilM, pTM = 0.84. D. AlphaFold3 model of TlaM, pTM = 0.47. E. AlphaFold3 model of the TslM-TlaM-TilM complex. Residues 1-25 of TilM not shown. Inset (*top*) the LFG motif in the TslM trafficking domain (blue), in predicted complex with TlaM (lilac). (*bottom*) Predicted Aligned Error plot of the complex generated by AlphaFold3, overall pTM score = 0.5, iPTM = 0.56.

The N-terminal 25 residues of the DUF1433 protein, TilM, are predicted to form an uncleaved transmembrane helix, with the remainder of the protein (aa 26 – 149) adopting a single folded domain (Fig 2A, Fig2C, Fig S6). TilM is substantially smaller than the TslA immunity protein, TilA, and the two immunity proteins do not share detectable sequence similarity, indicating that they represent different protein families. The other conserved protein encoded at the *tslM* locus, TlaM, is a 99-residue protein with a likely helical hairpin fold characteristic of the WXG100 family (Fig 2D, Fig S6). As expected, AlphaFold3 strongly predicts both TilM and TlaM to interact with TslM (Fig 2E). TilM is predicted to interact with the N-terminal lipase domain of TslM, while TlaM is predicted to associate with the C-terminal LXG domain of TslM, forming a helical bundle (Fig 2E). Within TslM’s LXG domain, a likely LFG secretion motif was identified due to both its conservation and positioning at the turn of the helices (Fig 2E inset).

### TslM is secreted by the T7SSb and requires TlaM

To determine whether TslM is secreted by the T7SS, we employed a split nanoluciferase assay previously used to investigate T7SS secretion (Garrett et al., 2023; Yang et al., 2024). Briefly, full length TslM was produced as a C-terminal fusion with the 11 amino acid pep86 fragment of nanoluciferase and expressed alone, or alongside TlaM, from the inducible expression plasmid pRAB11 in wild type *S. aureus* USA300 or an otherwise isogenic Δ*essC* mutant (Garrett et al., 2023). Following supplementation of the culture supernatant with the nanoluciferase large subunit, 11S, and furimizine substrate, we observed substantial luminescence from the strain co-producing TslM and TlaM, but not TslM only (Fig 3A). This indicates that TslM is secreted, but that this requires presence of the Lap-like partner protein TlaM. This is in line with observations showing that secretion of other T7SSb toxins is dependent on binding of Lap partners (Garrett et al., 2023; Klein et al., 2022, 2024; Yang et al., 2023). Secretion of TslM is strictly dependent on an active T7SS as no significant secretion was observed in the *essC* mutant strain (Fig 3A), with TslM accumulating in the cytoplasmic fraction in this strain background (Fig S8). As expected, only the LXG-like ‘trafficking’ domain of TslM is required for secretion as co-production of a pep86 fusion to just this domain alongside TlaM resulted in significant levels of nanoluciferase activity. We conclude that TslM is a substrate of the *S. aureus* T7SS and that it harbours a C-terminal secretion domain.

**Figure 3.**
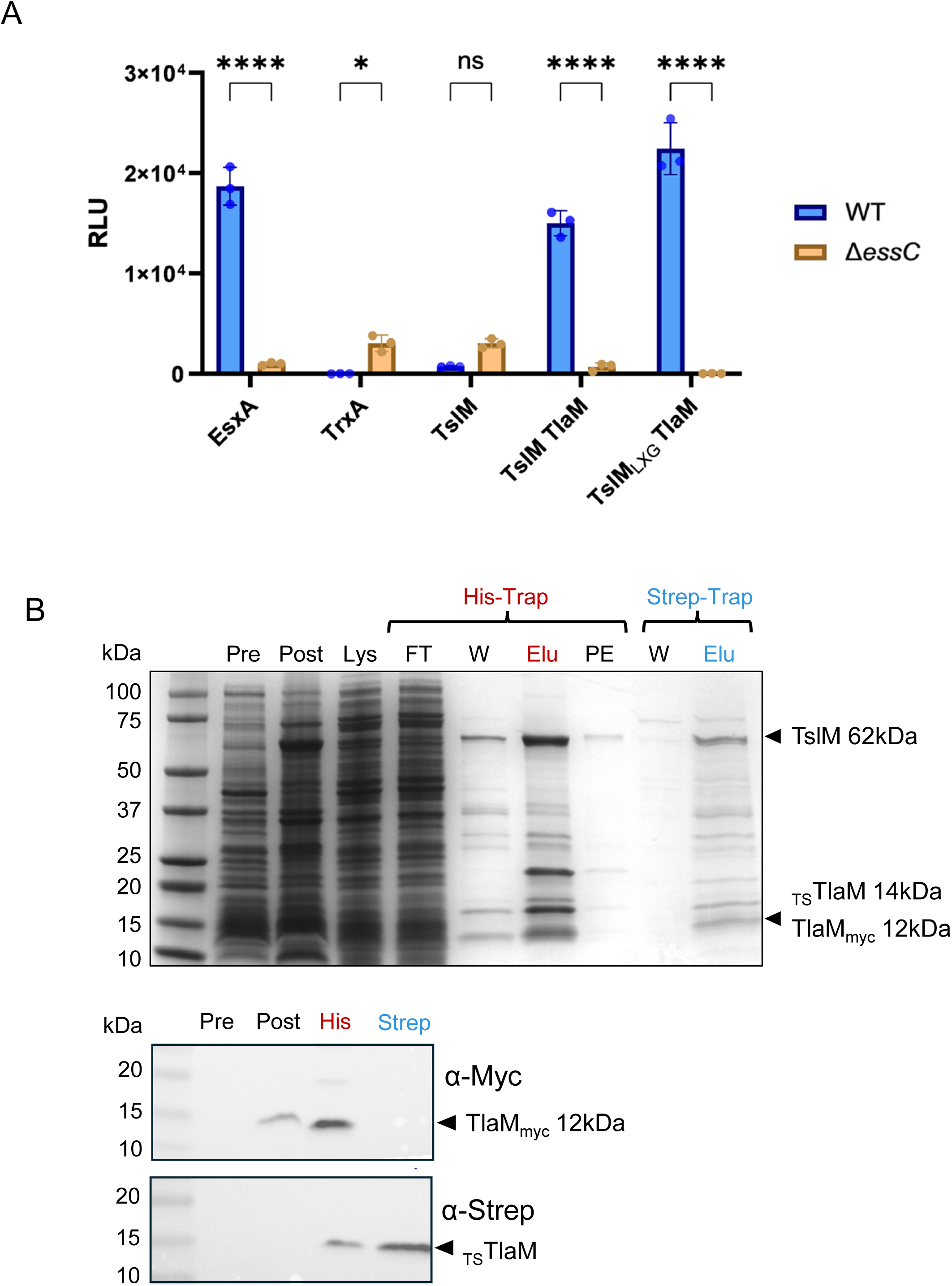
TslM is secreted by the *S. aureus* T7SS in a TlaM-dependent manner. A. Relative luminescence of supernatant fraction from cultures producing C-terminal pep86-tagged TslM alone, or with TlaM, or a pep86-tagged C-terminal domain alongside TlaM (TslM_LXG_-TlaM) in both wild type (blue) and Δ*essC* (orange) strains. C-terminally pep86-tagged EsxA (T7SS-secreted substrate) and TrxA (cytoplasmic protein) were included as controls. Readings were taken at maximum luminescence which reached 7 minutes following the addition of luciferase substrate and 11S. Error bars show standard deviation of three biological replicates. Significant difference was determined by 2way ANOVA: P values represented by asterisks: * <0.05, **** <0.0001. The corresponding cytoplasmic luminescence readings can be found in Fig S8. RLU – relative luminescence units. B. Co-production of TslM with two differentially-tagged TlaM proteins shows binding of only a single TlaM partner. Lysates of *E. coli* co-producing TslM-His alongside TlaM-Myc and TwinStrep-TlaM were subjected to either IMAC or StrepTrap chromatography. Fractions were analysed by SDS PAGE (top panel) or western blotting with either α-Myc (middle panel) or α-Strep (bottom panel) as indicated. Pre: pre-induction sample; Post: 4 hours post induction sample; Lys: lysis sample; FT: His column flow-through; W: wash sample; Elu: elution samples.

### TslM forms a complex with a single TlaM partner

Other characterised toxins usually have two distinct Lap partners (e.g. (Alcock et al., 2025; Garrett et al., 2023; Klein et al., 2022, 2024; Yang et al., 2024)), while EsaD has three (Yang et al., 2023). However, TslM is unusual as it is encoded next to a single partner, TlaM. We therefore wondered if a single copy of TlaM would bind to TslM, or whether it might bind as a dimer (or even higher order multimer). To test this, a vector was constructed encoding two copies of TlaM, differentially tagged with either Myc or TwinStrep and expressed alongside full length TslM with a C-terminal His tag. Complexes with isolated by either IMAC or streptavidin affinity chromatography in parallel, to pull down complexes containing either TslM-His or TlaM-TwinStrep. Eluted fractions from both purifications were then blotted with α-Myc and α-TwinStrep antibodies. As expected, the TslM-containing fractions from IMAC also contained both TwinStrep- and Myc-tagged TlaM, as TslM could complex with either tagged partner (Fig 3B). However, isolation of TlaM-TwinStrep using streptavidin affinity chromatography resulted in co-purification of TslM-His, but not Myc-tagged TlaM, indicating that complexes containing two different copies of TlaM do not exist (Fig 3B).

### Secreted TslM is toxic to *S. aureus* in the absence of TilM

We next tested whether TslM was toxic to *S. aureus*. To this end, we constructed a strain of USA300 deleted for all four Til2 (DUF1433)-encoding genes (*NRS_RS13750*, *NRS_RS13755* and *NRS_RS13765* at the *lplIII* locus and *NRS_RS09835* at the νSaβ island). When TslM and TlaM were expressed in both wild type and *til2* deletion strains, a small, but repeatable reduction in optical density was observed during growth (Fig 4A) and a significant reduction in colony counts recovered after 8 hours (Fig 4B). Furthermore, inactivation of the T7SS via deletion of *essC* resulted in the restoration of normal growth, indicating that the toxin must be secreted by the T7SS to exert its deleterious effect (Fig 4A,B).

**Figure 4.**
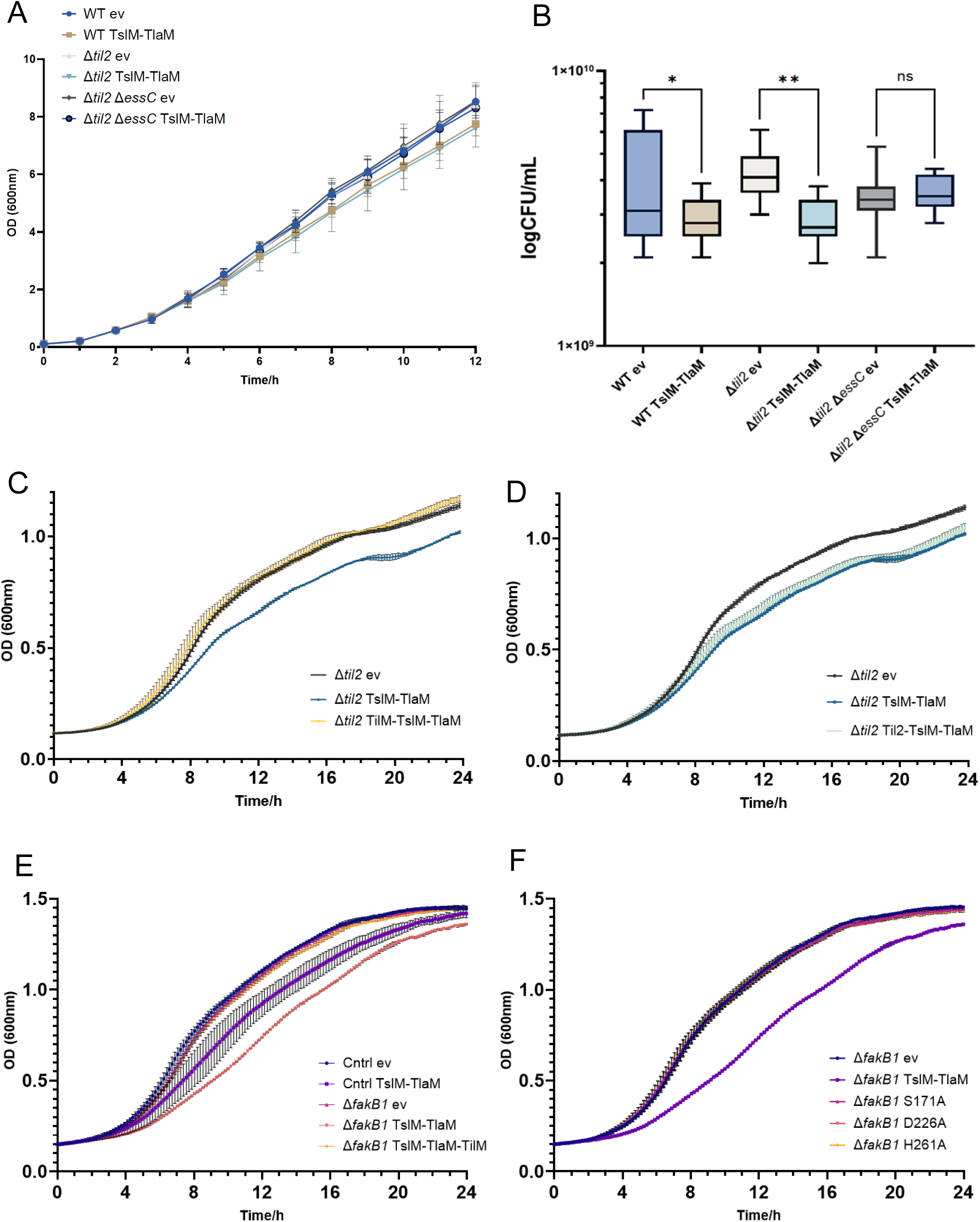
TslM is toxic to *S. aureus* and TilM protects against toxicity. A. Growth of either *S. aureus* USA300, USA300/Δ*til2* or USA300/Δ*til2/*Δ*essC* harbouring either empty pRAB11 (ev) or pRAB11-TslM-TlaM as indicated. Cultures were supplemented with ATc at 2 hours to induce plasmid-encoded gene expression. Error bars show standard deviation of three biological replicates. B. Log(colony forming units)/mL of the cultures assessed after eight hours growth (n = 3). Significant difference was determined by 2way ANOVA: P values represented by asterisks: * <0.05, ** <0.01. C. Growth of USA300/Δ*til2* harbouring empty pRAB11 (black), pRAB11-TslM-TlaM (blue) or pRAB11-TslM-TlaM-TilM (yellow), (n = 3 biological replicates). D. Growth of Δ*til2* strains expressing empty pRAB11 (black), pRAB11-TslM-TlaM (blue) or pRAB11-TslM-TlaM-Til2 (sea green), (n = 3). “Til2” here refers to NRS_RS09835, a non-cognate DUF1433 encoded in *S. aureus* USA300. E. Growth of Δ*fakB1* strains from the NTML library (strain NE1540) expressing empty pRAB11 (pink), pRAB11-TslM-TlaM (salmon) or pRAB11-TslM-TlaM-TilM (yellow), (n = 3). Controls (Cntrl) are strain NE1612 from the NTML library, with a transposon insertion into a pseudogene, expressing empty pRAB11 (blue) or pRAB11-TslM-TlaM (violet). F. Growth of Δ*fakB1* strains from the NTML library expressing empty pRAB11 (blue), pRAB11-TslM-TlaM (violet), or pRAB11-TslM-TlaM with a catalytic substitution: S171A (pink), D226A (salmon), H261A (yellow), (n = 3).

To probe the toxicity of TslM further, we adapted our growth analysis to 96-well plate format. Production of TslM was associated with a significant reduction in growth, in both the wild type and the *til2* deletion strains. Co-production of the immunity protein, TilM, in either strain background was sufficient to fully alleviate the toxicity associated with TslM production (Fig 4C). The toxicity of TslM to the wild type USA300 strain was surprising given that it encodes four Til2-family proteins. However, none of these are identical to TilM (Fig S9). We considered two explanations for the lack of protection from TslM toxicity – either the chromosomally-encoded Til2 proteins are not produced at sufficient levels to neutralise TslM, or they do not bind sufficiently tightly to TslM to block its activity. To distinguish these, we replaced the gene coding for TilM on the *tilM-tslM-tlaM* expression construct with that of NRS_RS09835, the only Til2 protein encoded at the USA300 νSaβ island. Expression of this construct resulted in a similar reduced growth phenotype as the construct lacking TilM (Fig 4D). We conclude that the susceptibility of wild type USA300 to TslM toxicity is because the Til2 homologues do not provide full protection against TslM from MRSA252. In support of this AlphaFold3 predictions suggest that the binding of NRS_RS09835 to the lipase domain of TslM is significantly weaker than the cognate interaction of TilM-TslM, and that it does not occlude the active site in the same way that TilM is predicted to do (Fig S10).

Since TslM is a predicted lipase, we wondered whether toxicity might be enhanced in a *fakB1* background. FakB1 is a fatty acid binding protein and deletion of the encoding gene results in impaired ability to repair membrane damage, but no growth defects or lipidomic abnormalities under standard conditions (Machinandiarena et al., 2020; Parsons et al., 2014). As shown in Fig 4E, TslM exhibited enhanced toxicity in the USA300 *fakB1* mutant strain background in the absence of TilM, consistent with TslM targeting membrane lipids. To confirm that the reduced growth phenotype in these experiments arose due to TslM activity, we constructed substitutions in the predicted active site of TslM: S171A, D226A and H261A. Each of these substitutions restored wild type growth to the USA300 Δ*fakB1* strain (Fig 4F).

### TslM intoxicated cell membranes accumulate free fatty acids

The results presented above are consistent with TslM having lipase activity. To confirm this, we isolated membranes from the *fakB1* mutant strain producing TslM, the non-toxic S171A variant, or wild type TslM alongside TilM, and carried out mass spectrometry lipidomics (Fig. 5). Membranes isolated from cells producing wild type TslM showed accumulation of *lyso*-PG (Fig 5, inset) and free fatty acid compared to membranes from strains harbouring an empty plasmid vector, or producing either TslM_S171A_ or wild type TslM alongside its TilM immunity protein. These findings are consistent with TslM being an anti-staphylococcal lipase that damages membranes of susceptible bacteria.

**Figure 5.**
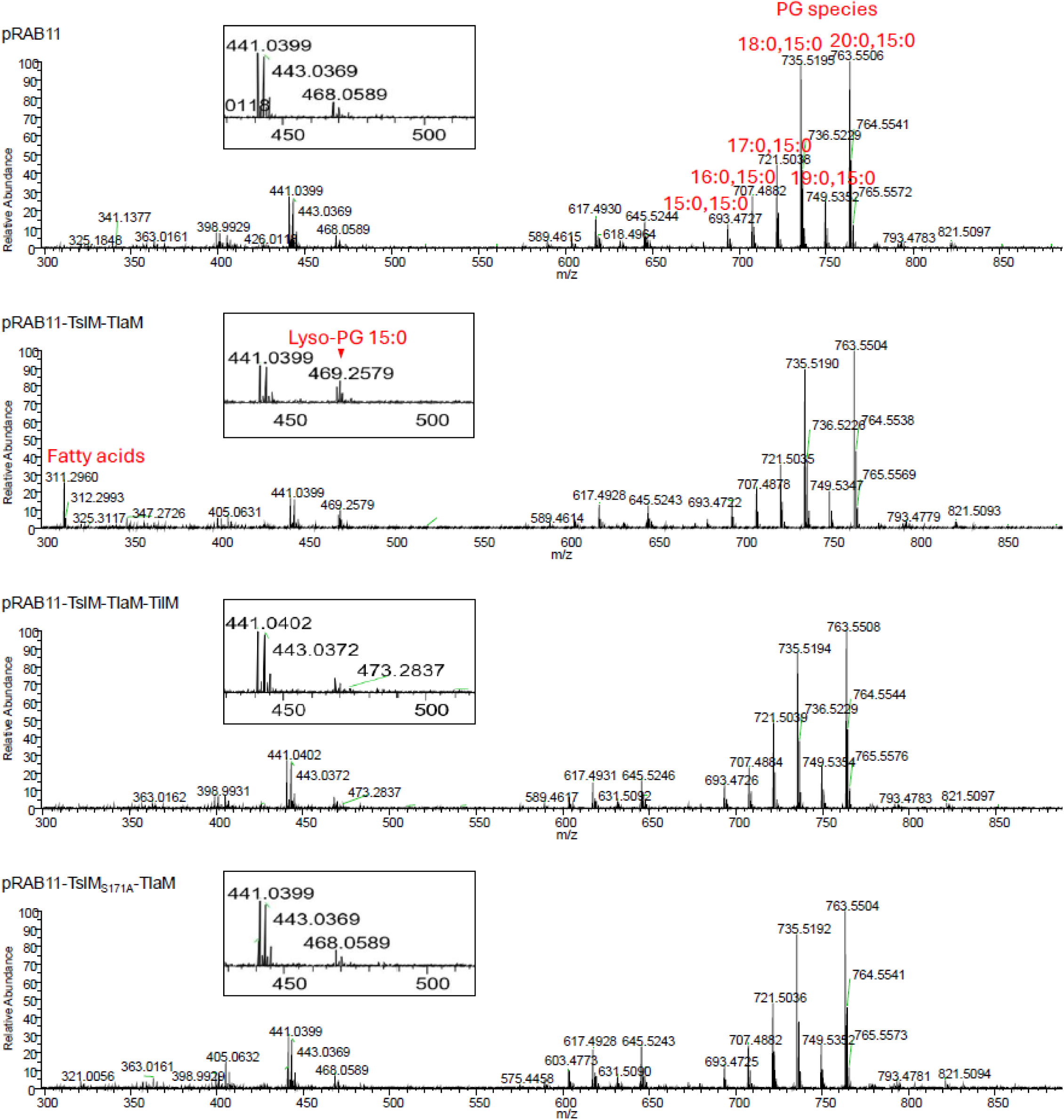
Lipidomic analysis of *S. aureus* membranes following TslM intoxication. Positive ion mode mass spectra of membrane lipids extracted the USA300/Δ*fakB1* strain harbouring either empty pRAB11, pRAB11-TslM-TlaM, pRAB11-TslM-TlaM-TilM, or pRAB11-TslM-TlaM with the TslM S171A catalytic substitution. Lipids were extracted at 6 hours post supplementation with ATC to induce plasmid-encoded gene expression. Identified phosphatidylglycerol (PG) species, *lyso*-PG and free fatty acids are labelled. Insets: magnified region between m/z 450-500.

## Discussion

In this study we have identified the Tsl2 family as novel substrates of the *S. aureus* T7SS. This is only the second family of ‘reverse’ T7SS secreted substrates to be identified, after Tsl1, and is also widespread, with homologues encoded in many Bacillota species. The two toxin families share overall similar structural arrangement with N-terminal lipase toxin domains, and C-terminal LXG-like helical domains that contain the T7SS-targeting information. However, there are also key differences - for example the immunity proteins are from different families and Til2 are transmembrane proteins while Til1 are lipoproteins. Moreover, TilA requires two Lap-like partners to mediate its secretion whereas TilM is unusual in requiring only a single Lap partner.

It is striking that while only two families of ‘reverse’ T7SS substrates have been identified to date, both are phospholipases. It is also noteworthy that no lipases have yet been identified with N-terminal LXG domains, although a lipid II phosphatase toxin with an N-terminal secretion domain is a substrate of the T7SSb of *S. intermedius* B196 (Klein et al., 2022). It is not clear whether phospholipases have features that require them to be fused N-terminal to the T7SS-targeting sequence, although it should be noted that other staphylococcal phospholipases are synthesised as pre-pro-proteins and are N-terminally processed in the cell envelope following their secretion (Götz et al., 1998). In future it would be interesting to determine whether fusion of the TslM or TslA lipase domains to a canonical LXG domain would permit the secretion of a functionally active toxin.

Previous work has shown that the Tsl1 family of reverse T7SS lipases can be found in up to three paralogous copies in *S. aureus* genomes (Garrett et al., 2023). Here we identified four Tsl2 paralogues, three of which are encoded on chromosomal islands. Aside from *tslM* which is found on the νSaβ island, *tslN* is present on the νSaα island and *tslP* is found on multiple SaPIs which can integrate into at least four different loci in the *S. aureus* chromosome. This suggests that *S. aureus* has the capacity to encode up to seven non-identical copies of Tsl2 proteins in their genomes. Taken together our findings show that the reverse lipase toxins are the most numerically abundant secreted toxins in *S. aureus* and are therefore likely to be important players in competitive environments. It remains to be seen whether other toxin domains may also be found in a reverse targeting arrangement.

## Methods

### Bacterial Strains and growth conditions

All *S. aureus* strains used in this study are listed in Table S1. *S. aureus* was grown in tryptic soy broth (TSB) or on TS agar at 37°C unless otherwise stated. Media was supplemented with chloramphenicol (Cml; 10 μg/mL) and erythromycin (Ery; 5 μg/mL) where necessary for plasmid and transposon maintenance, respectively. Induction of plasmid-encoded gene expression was achieved through supplementation of growth media with anhydrotetracycline (ATc, 500 ng/mL), and counterselection during allelic exchange with pIMAY using 100 ng/mL ATc. *Escherichia coli* strains used in this study can be found in Table S1. *E. coli* was grown in lysogeny broth (LB) or on LB agar at 37°C unless otherwise stated. Media was supplemented with ampicillin (Amp; 100 μg/mL), chloramphenicol (25 μg/mL) and kanamycin (Kan; 50 μg/mL) where necessary for plasmid maintenance. Induction of plasmid-encoded gene expression was achieved through supplementation of Isopropyl β-D-1-thiogalactopyranoside (IPTG, 500 μM).

For growth curve analysis, *S. aureus* cultures were supplemented with Cml and 2 mM CaCl_2_, (as lipases can depend on Ca^2+^ for activity (Rosenstein & Götz, 2000)), and plasmid-encoded gene expression was induced with ATc at 2 hours. OD_600_ readings were taken manually every hour, in triplicate, over 12 hours. Each experiment was performed with three biological replicates. Colony forming units (CFU) were enumerated by serial dilution of cultures taken from growth curve experiments every four hours (0, 4, 8, 12), followed by plating. Counts were calculated with five technical replicates per strain and three biological replicates per experiment. For growth analysis using a plate reader, *S. aureus* cultures were grown overnight and then normalised to OD_600_ 0.1 in fresh TSB containing Cml or Cml and Ery, whereupon they were supplemented with 2 mM CaCl_2_ and 500 ng/mL ATc. Strains were then added, in triplicate, to wells of a 96-well clear plate (Greiner) at a volume of 150 µL per well and grown in a Tecan Infinite Nano M+ plate reader at 30°C, shaking at 1.5mm amplitude. OD_600_ was measured every 7 minutes for 24 hours. Each experiment was performed with three biological replicates.

### Plasmid and strain construction

Plasmids were constructed by Gibson assembly. Plasmids used in this study are listed in Table S2. Catalytic substitutions were introduced using PCR amplification followed by ligation with T4 ligase and polynucleotide kinase. All plasmids were verified with whole plasmid sequencing carried out by Plasmidsaurus (https://plasmidsaurus.com/). Constructs in pQE70 for the recombinant expression of TslM, TlaM and TilM were constructed with the fusion of C-terminal His6 (-HHHHHH*), C-terminal Myc (-EQKLISEEDL*) or N-terminal Twin-Strep tags (MGSGWSHPQFEKGGGSG GGSGGGSWSHPQFEKGSG-), respectively. TslM, TlaM and TilM sequences were amplified from MRSA252 genomic DNA before assembly in *E. coli. S. aureus* deletion strains were constructed using allelic exchange with either the pIMAY vector, which was used to construct the Δ*til2* Δ*essC* strain, or otherwise the pIMAYZ vector (Monk & Stinear, 2020). Briefly, plasmids were designed with 500bp upper and lower arms corresponding to upstream and downstream regions of the gene to be deleted, including the first and last six codons as a scar region to be left in place. Up- and downstream regions were amplified from *S. aureus* USA300 genomic DNA and assembled in *E. coli*, then electroporated into *S. aureus.* Following integration and excision steps as described in the literature, deletion mutants were verified by PCR amplification of the region of interest and then whole genome sequencing, carried out by Plasmidsaurus.

### Nanoluciferase secretion assay

This was performed essentially as described (Garrett et al., 2023; Yang et al., 2024). Strains were grown overnight and subcultured into fresh TSB containing Cml to a starting OD_600_ of 0.1. Cells were grown for 2 hours before induction of pep86 fusion production by addition of ATc. Cells were grown for a further 2 hours to an OD_600_ of ∼2. To obtain the cytoplasmic/supernatant fractions, the equivalent of 1 mL of sample of OD_600_ = 1 was withdrawn for each sample and pelleted by centrifugation. The top 150 μL of supernatant following pelleting was withdrawn and kept as the supernatant fraction. The cell pellet was resuspended in 500 µL Tris buffered saline (TBS; 20 mM Tris pH 7.6, 137 mM NaCl) supplemented with 100 µg/mL lysostaphin and incubated at 37°C for 45 minutes. The cell samples were then boiled for 10 minutes to fully lyse cells and pelleted at full speed for 30 seconds to remove cell debris. Samples were cooled to room temperature and serially diluted three times 1 in 2, in TBS. 150 μL of each dilution, in triplicate, were aliquoted into a Greiner CELLSTAR white 96 well plate. Furimazine solution (Promega Cat. #N1610) was diluted 1/100 in TSB. To each well, 10 µL of 5:2 purified 11S (purified as described in Garrett et al., 2023):furimazine solution was added and the luminescence read at 1 minute intervals for 10 minutes, using the FLUOstar Omega reader with a gain value of 3,000. Data were analysed to find the peak luminescence signal and this was used to visualise the data.

### Recombinant protein purification

6 L cultures of terrific broth (Formedium) supplemented with Amp/Kan and 24 mL glycerol were inoculated from an overnight culture (at 1/100 dilution) of *E. coli* M15 (pREP4) carrying the appropriate recombinant protein expression construct. After 2 hours growth, 500 μM IPTG was added to induce plasmid-encoded gene expression. After a further 4 hours, cells were harvested by centrifugation and pellets resuspended in 5 mL lysis buffer per gram (lysis buffer: 25 mM imidazole, 50 mM Tris-HCl pH 7.5, 500 mM NaCl (or 150 mM NaCl for two partner TlaM experiment)). Cells were lysed by passage through an emulsifier at 18000 psi and the lysate clarified by centrifugation. Tagged proteins were then purified from clarified lysate by immobilised metal (nickel) ion affinity chromatography (IMAC) on an ÄKTA platform, using an imidazole gradient (elution buffer: 500 mM imidazole, 50 mM Tris-HCl pH 7.5, 500 mM NaCl (or 150 mM NaCl for two partner TlaM experiment)). For StrepTactin purification, proteins were lysed in 50 mM HEPES pH 7.8, 150 mM NaCl, 1 mM EDTA and then purified by StrepTactin affinity chromatography on an ÄKTA platform and eluted with biotin (elution buffer: BXT elution buffer 10x, IBA-Lifesciences – diluted 1/10 with StrepTactin lysis buffer). Protein fractions were resuspended in 4x Laemmli sample buffer with 2-mercaptoethanol and boiled for 15 minutes to denature proteins before further analysis.

### SDS-PAGE and western blot analysis

SDS- PAGE gels were electrophoresed at 80 V for 10 minutes followed by 200 V for 35 minutes, and protein visualised by staining with Coomassie Instant Blue. For western blot, proteins were transferred onto nitrocellulose membrane in transfer buffer (25 mM Tris, 192 mM glycine pH 8.3, 20% ethanol) at 25 mA for 90 minutes. Western blotting was performed against His_6_, Twin-Strep and Myc tags using the following antibodies: HRP-linked α-His (Sigma A7058-1VL), HRP-linked α-Strep (IBA-Lifesciences 2-1509-001) and HRP-linked α-Myc (abcam ab1326). 5% milk in TBS-T (Tris buffered saline plus 0.5% Tween 20) was used as a blocking agent.

### Lipid extractions and mass spectrometry

*S. aureus* overnight cultures were subcultured to OD_600_ 0.1 in fresh TSB supplemented with Cml/Ery and 2 mM CaCl_2_ and grown for 8 hours. After 2hours, ATc was added to induce plasmid-encoded gene expression. At 8 hours, cell lipids were extracted according to the method of Bligh and Dyer (1959) as previously reported (Garrett et al., 2023). Briefly, cells were washed with PBS, re-suspended in 100 µL of PBS and transferred to a glass vial, then 375 µL of 2:1 (v/v) methanol:chloroform was added. This was vortexed and agitated for at least 10 minutes, after which 125 µL of water and 125 µL of chloroform was added, and the mixture agitated to make it biphasic. After settling for 30 minutes, the lower organic, chloroform-rich phase was removed and placed into a fresh glass vial. This organic phase was dried before further analysis. Mass spectrometry was carried out as reported previously (Garrett et al., 2023). Organic phases were suspended in 2:1 (v/v) methanol:chloroform and high-resolution mass spectrometry data were acquired by electrospray ionisation techniques using a Thermo Scientific^TM^ Exactive^TM^ Orbitrap mass spectrometer. Phospholipid species annotations were determined in reference to previous assignments and the LIPID MAPS database (https://www.lipidmaps.org).

### Bioinformatic analysis

BLASTp searches were performed against the RefSeq database (Johnson et al., 2008; Pruitt et al., 2005) to find homologues of DUF1433 and Tsl2 proteins. Flanking gene analysis of accession lists was run through FlaGs and the webFlaGs server with default settings (Saha et al., 2021). Clinker was used to visualise and compare loci (Gilchrist & Chooi, 2021). Protein sequences were aligned with the MUSCLE algorithm in Jalview (Procter et al., 2021) for multiple sequence alignments, or the EMBOSS Water tool (Smith-Waterman algorithm) for pairwise sequence alignment. All alignments were visualised in Jalview and coloured by sequence conservation. Maximum likelihood tree of DUF1433 protein sequences was generated in Jalview. Protein structural models were generated, unless otherwise stated, using the AlphaFold3 server (Abramson et al., 2024). Structural models were visualised and rendered in CCP4MG (McNicholas et al., 2011). Superpositions were achieved using the SSM method in CCP4MG.

## Acknowledgements

This study was supported by the Wellcome Trust (through Investigator Awards 10183/Z/15/Z and 224151/Z/21/Z to TP). ABH was funded by the Newcastle-Liverpool-Durham BBSRC DTP3 Training Grant, project reference number BB/T008695/1 and SRG was funded by the Newcastle-Liverpool-Durham BBSRC DTP2 Training Grant, project reference number BB/M011186/1. We would like to thank Dr Kieran Bowran for his help with running large scale FlaGs anaylsis.

## Author Contributions

ABH, SG, TKS, TRB and TP designed experiments. ABH, SRG, JS and TKS carried out experimental work. ABH, SRG, JS, TKS and TP undertook data analysis. TP and ABH wrote the manuscript which was edited by all other authors. All authors have approved the final version.

## Competing Interests

the authors declare no competing interests.

## Notes

### Competing Interest Statement

The authors have declared no competing interest.

