## supplementary Tables and Figures for "A novel reverse lipase toxin substrate of the *Staphylococcus aureus* type VII secretion system"

SUPPLEMENTARY INFORMATION

**Table S1. Bacterial strains used in this study**

| Strain | Genotype/description | Source/Reference |
|---|---|---|
| *E. coli* | | |
| DH5α | φ80d Δ(*lacZ*)M15 *recA1 endA1 gyrA96 thi-1 hsdR17* (rk-mk+) *supE44 relA1 deoR* Δ(*lacZYA-argF*)U169 | Promega |
| JM110 | *rpsL thr leu thi lacY galK galT ara tonA tsx dam dcm glnV44* Δ(*lac-proAB*) e14- [F' *traD36 proAB*+ *lacIq lacZ*ΔM15] *hsdR17*(rK−mK+) | Stratagene |
| M15 (pREP4) | F-, *lac*, *ara*, *gal*, *mtl*, [Kan$^r$, *lacI*] | Qiagen |
| *S. aureus* | | |
| MRSA252 | Wild type | (Holden et al., 2004) |
| USA300 LAC | Wild type | (Kazakova et al., 2005) |
| USA300 Δ*essC* | In-frame deletion of *essC* from USA300 LAC | (Garrett et al., 2023) |
| USA300 Δ*til2* | In-frame deletion of all four *DUF1433* (*til2*) genes in USA300 LAC: three at the *lpIII* locus and one at the vSaβ island | This work |
| USA300 Δ*til2* Δ*essC* | As above, also Δ*essC* | This work |
| USA300 JE2 NE1540 | Tn insertion [Ery$^r$] in *fakB1* (*SAUSA300_0733*) | NTML (Fey et al., 2013) |
| USA300 JE2 NE1612 | Tn insertion [Ery$^r$] in prophage tail component (*SAUSA300_1397*) | NTML (Fey et al., 2013) |

**Table S2. Plasmids used in this study**

| Plasmid | Description | Source/Reference |
|---|---|---|
| pRAB11 | *E. coli*/*S. aureus* shuttle vector, inducible expression, $Amp^r$, $Cml^r$ | (Helle et al., 2011) |
| pRAB11-$TslM_{pep86}$ | pRAB11 encoding TslMpep86 fusion, preceded by *hla* rbs | This work |
| pRAB11-$TslM_{pep86}$-TlaM | As pRAB11-$TslM_{pep86}$ with TlaM also encoded | This work |
| pRAB11-$TslMLXG_{pep86}$-TlaM | As pRAB11-$TslM_{pep86}$-TlaM with only the LXG domain of TslM encoded | This work |
| pRAB11-$EsxA_{pep86}$ | pRAB11 encoding EsxA with C-terminal pep86 fusion | (Yang et al., 2024) |
| pRAB11-$TrxA_{pep86}$ | pRAB11 encoding TrxA with C-terminal pep86 fusion | (Yang et al., 2024) |
| pQE70 | Vector for *E. coli*, inducible expression, T5 pro, $Amp^r$ | Qiagen |
| pQE70-$TslM_{His6}$-$_{TS}$TlaM-$TlaM_{Myc}$ | pQE70 encoding TslM with a C-terminal His6 fusion and two copies of TlaM with an N-terminal TwinStrep fusion and C-terminal Myc fusion | This work |
| pIMAYZ | *E. coli*/*S. aureus* shuttle vector, temperature sensitive, $Cml^r$ | (Monk & Stinear, 2020) |
| pIMAYZ-til2_lplIII | pIMAYZ carrying *til2* deletion allele for the *lplIII* locus | This work |
| pIMAYZ-til2_vSaβ | pIMAYZ carrying *til2* deletion allele for the vSaβ island | This work |
| pIMAY-essC | pIMAY carrying *essC* deletion allele | (Garrett et al., 2023) |
| pRAB11-TslM-TlaM | pRAB11 encoding TslM-TlaM, preceded by *hla* rbs | This work |
| pRAB11-TilM-TslM-TlaM | As pRAB11-TslM-TlaM with TilM (full length) | This work |
| pRAB11-Til2-TslM-TlaM | As pRAB11-TslM-TlaM with NRS_RS09835 (full length) | This work |
| pRAB11-$TslM_{S171A}$-TlaM | As pRAB11-TslM-TlaM with a S171A substitution in TslM | This work |
| pRAB11-$TslM_{D226A}$-TlaM | As pRAB11-TslM-TlaM with a D226A substitution in TslM | This work |
| pRAB11-$TslM_{H261A}$-TlaM | As pRAB11-TslM-TlaM with a H261A substitution in TslM | This work |

**Supplementary figure legends**

Figure S1. Gene neighbourhood analysis of DUF1433-encoding genes. The DUF1433 protein WP_001034756.1 was used in a webFlags search of *S. aureus* genomes present in RefSeq to identify where homologous proteins were encoded. From the output, genes shaded black or red (numbered 1) correspond to DUF1433 (Til2)-encoding genes. Genes shaded blue and numbered 3 correspond to *til1* (encoding immunity proteins to TslB), genes shaded orange and numbered 25 correspond to *tslB,* genes shaded pink and numbered 19 encode the Tsl2 family of reverse phospholipases (TslO in this instance) and genes shaded purple and numbered 20 encode Tla2 proteins (TlaO in this instance). White genes with a blue outline are pseudogenes.

Figure S2. Alignment of the phospholipase domains of TslO (WP_033856953.1) and TslA. Pairwise alignment performed with Smith-Waterman algorithm, visualised in Jalview. Blue residues/logo indicate residue matches. Phospholipase catalytic triad of TslA (S164, D224, H251) and aligning predicted catalytic triad of TslO shown with red arrows.

Figure S3. Gene neighbourhood analysis of novel phospholipase-encoding genes. The protein sequence of WP_033856953.1, a putative phospholipase, was used in a FlaGs search of *S. aureus* genomes present in RefSeq to identify where homologous proteins were encoded. From the output, genes shaded black correspond to Til2 lipase-encoding genes. The TslM, TslN, TslO and TslP-encoding families are indicated, alongside their chromosomal loci.

Figure S4. A. Protein sequence identity matrix of the four representative members of the Tsl2 family. Seq. ID was calculated by pairwise alignment. B. Maximum likelihood phylogenetic tree of the 305 Tsl2 protein sequences identified by BLAST search, as well as TslA. Families are coloured as follows: TslM (blue), TslO (purple), TslN/P (red), TslA (cyan). TslA groups with the smaller homologues, likely due to its relative size.TslP and TslN have high sequence similarity and are distinguishable by their locus, with *tslP* being encoded on SaPIs.

Figure S5. Alignment of vSaβ island subtypes that encode *tslM*. vSaβ genomic island subtypes III, VI, VIII, IX and XVII showing *tslM* (light blue) alongside *tilM* and *tlaM.* Also shown is the type II island from USA300, which does not encode *tslM* but still encodes an “orphan” Til2/DUF1433 (yellow). The majority of other vSaβ subtypes encode at least one Til2 (not shown but see (Kläui et al., 2019)). Alignment generated using Clinker.

Figure S6. Alphafold3 models of TslM, TilM and TlaM coloured by confidence (where red is low confidence and blue high confidence). Predicted Template Modeling scores (pTM) are also given.

Figure S7. Lip2 and TslM superposition. Superposition of the lipase domain of TslM (residues 1-268, blue) and Lip2, a characterised lipase (PDB ID: 3o0d(Bordes et al., 2010), grey). The predicted catalytic triad of TslM (S171A, D226A, H261A) is shown in red, and the catalytic triad of Lip2 (S162, D230, H289) in gold. Inset: close up view of the predicted active site of TslM superposed with that of Lip2, with surface view shown. Superposition was performed with the SSM method.

Figure S8. Nanoluciferase assay of cytoplasmic fraction of samples from Figure 4. Relative luminescence of cytoplasmic fraction from cultures producing C-terminal pep86-tagged TslM alone, or with TlaM, or a pep86-tagged C-terminal domain alongside TlaM ($TslM_{LXG}$-TlaM) in both wild type (blue) and Δ*essC* (orange) strains. C-terminally pep86-tagged EsxA (T7SS-secreted substrate) and TrxA (cytoplasmic protein) were included as controls. Readings were taken at maximum luminescence which reached 7 minutes following the addition of luciferase substrate and 11S. Error bars show standard deviation of three biological replicates. Significant difference was determined by 2way ANOVA: P values represented by asterisks: * <0.05, ** <0.01.

Figure S9. Multiple sequence alignment of the Til2 proteins encoded by USA300 and MRSA252. A. Alignment performed with the Muscle algorithm, visualised in Jalview. Residues and logo coloured by sequence conservation. *til2* genes are numbered from the first gene in a downstream direction in their respective loci: vSaβ or *lpl*III. Pseudogenes (indicated) were repaired for the purposes of the alignment by the manual removal of frameshifts and subsequent translation to the shown amino acid sequence. The conserved N-terminal transmembrane helix is boxed in red. Sequences are coloured and grouped according to the tree shown in B. B. Maximum likelihood phylogeny generated from the 15 Til2 proteins encoded in USA300 and MRSA252. Three clear clades emerge, with TilM grouping best with the other Til2 proteins encoded at the MRSA252 vSaβ locus, and all *lpl*III encoded Til2 proteins from both strains grouping together. This indicates that immunity protein diversity is locus specific, possibly reflecting the specific evolution of immunities required to protect against toxin subtypes within each toxin family. NRS_RS09835, used as the noncognate Til2 in toxicity experiments of Fig 7B, is labelled here as USAvSaβ.

Figure S10. Modelling of Til2 binding to TslM. A. TslM lipase domain (blue) and TilM (without TM helix, gold) complex generated by AlphaFold3. Inset: Predicted Aligned Error plot of the complex generated by AlphaFold3, iPTM = 0.89. B. TslM lipase domain (blue) and noncognate Til2 NRS_RS09835 (without TM helix, orange) complex generated by AlphaFold3. Inset: Predicted Aligned Error plot of the complex generated by AlphaFold3, iPTM = 0.23. Note the difference in predicted binding position and ipTM confidence scores between the complexes.

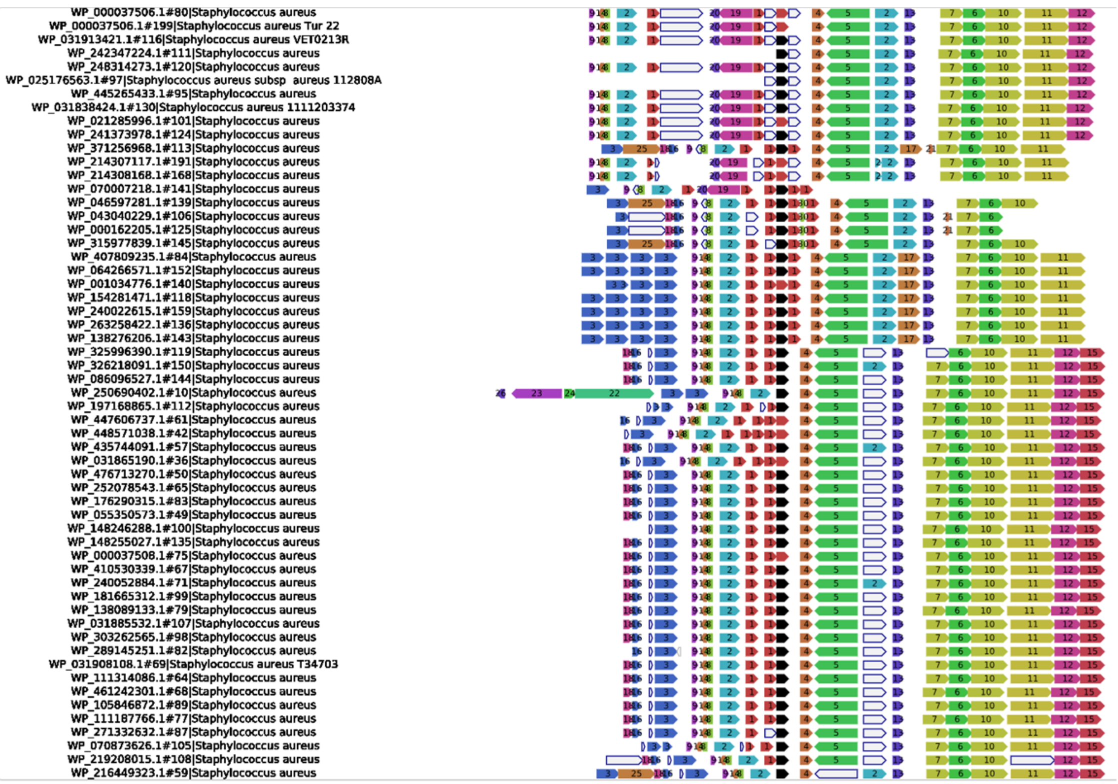


**Figure S1.** Gene neighbourhood analysis of DUF1433-encoding genes. The DUF1433 protein WP_001034756.1 was used in a webFlags search of *S. aureus* genomes present in RefSeq to identify where homologous proteins were encoded. From the output, genes shaded black or red (numbered 1) correspond to DUF1433 (Til2)-encoding genes. Genes shaded blue and numbered 3 correspond to *til1* (encoding immunity proteins to TslB), genes shaded orange and numbered 25 correspond to *tslB,* genes shaded pink and numbered 19 encode the Tsl2 family of reverse phospholipases (TslO in this instance) and genes shaded purple and numbered 20 encode Tla2 proteins (TlaO in this instance). White genes with a blue outline are pseudogenes.

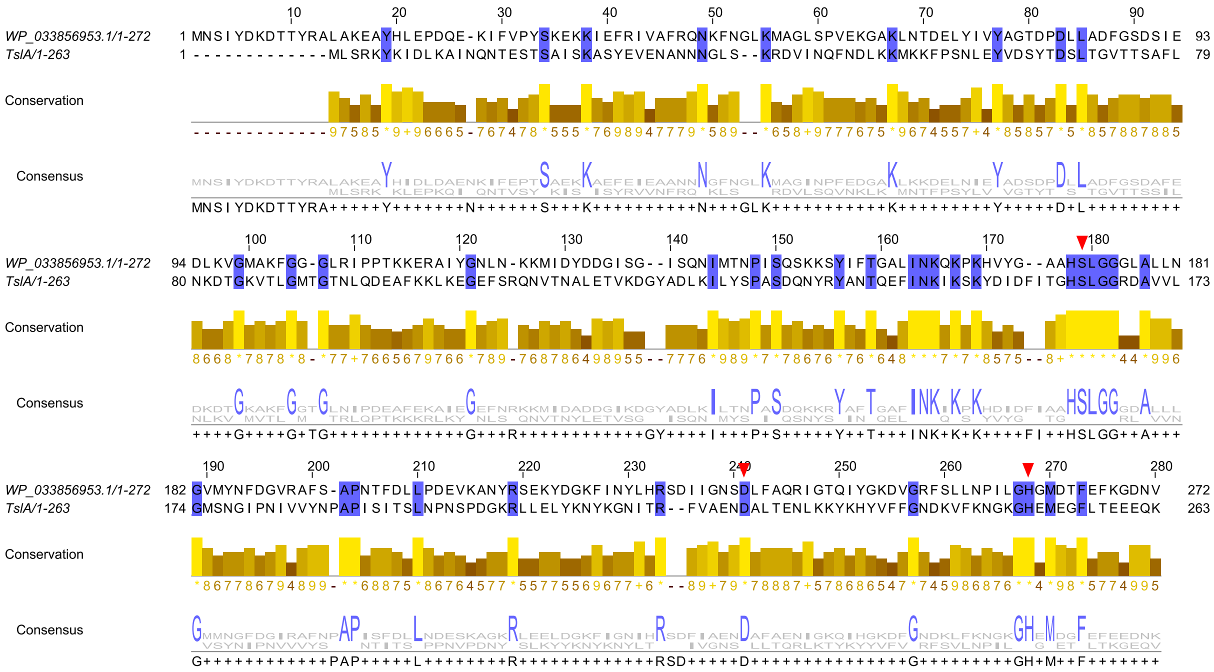

**Figure S2.** Alignment of the phospholipase domains of TslO (WP_033856953.1) and TslA. Pairwise alignment performed with Smith-Waterman algorithm, visualised in Jalview. Blue residues/logo indicate residue matches. Phospholipase catalytic triad of TslA (S164, D224, H251) and aligning predicted catalytic triad of TslO shown with red arrows.

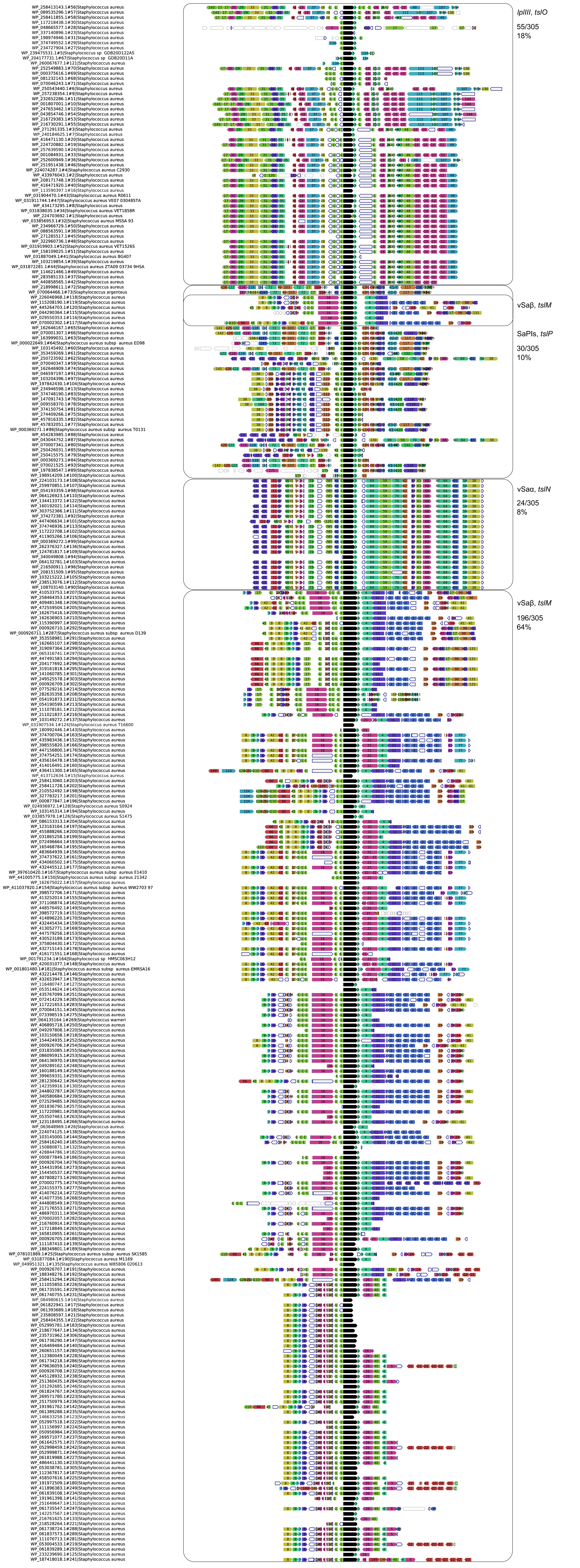
lpIIII, tslO
55/305
18%
vSaβ, tslM
SaPIs, tslP
30/305
10%
vSaα, tslN
24/305
8%
vSaβ, tslM
196/305
64%

**Figure S3.** Gene neighbourhood analysis of novel phospholipase-encoding genes. The protein sequence of WP_033856953.1, a putative phospholipase, was used in a FlaGs search of *S. aureus* genomes present in RefSeq to identify where homologous proteins were encoded. From the output, genes shaded black correspond to Til2 lipase-encoding genes. The TslM, TslN, TslO and TslP-encoding families are indicated, alongside their chromosomal loci.

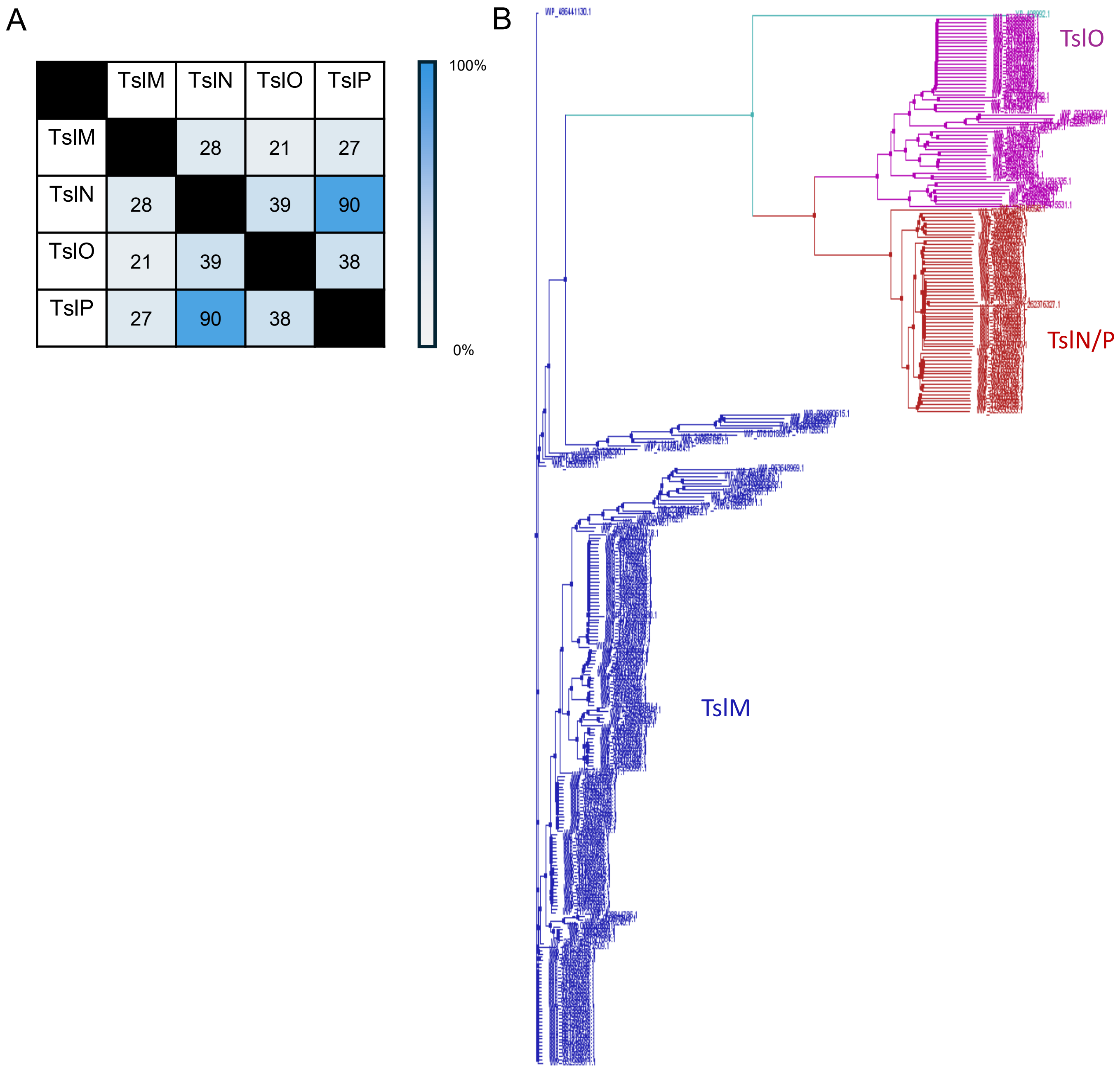


**Figure S4.** A. Protein sequence identity matrix of the four representative members of the Tsl2 family. Seq. ID was calculated by pairwise alignment. B. Maximum likelihood phylogenetic tree of the 305 Tsl2 protein sequences identified by BLAST search, as well as TslA. Families are coloured as follows: TslM (blue), TslO (purple), TslN/P (red), TslA (cyan). TslA groups with the smaller homologues, likely due to its relative size.TslP and TslN have high sequence similarity and are distinguishable by their locus, with *tslP* being encoded on SaPIs.

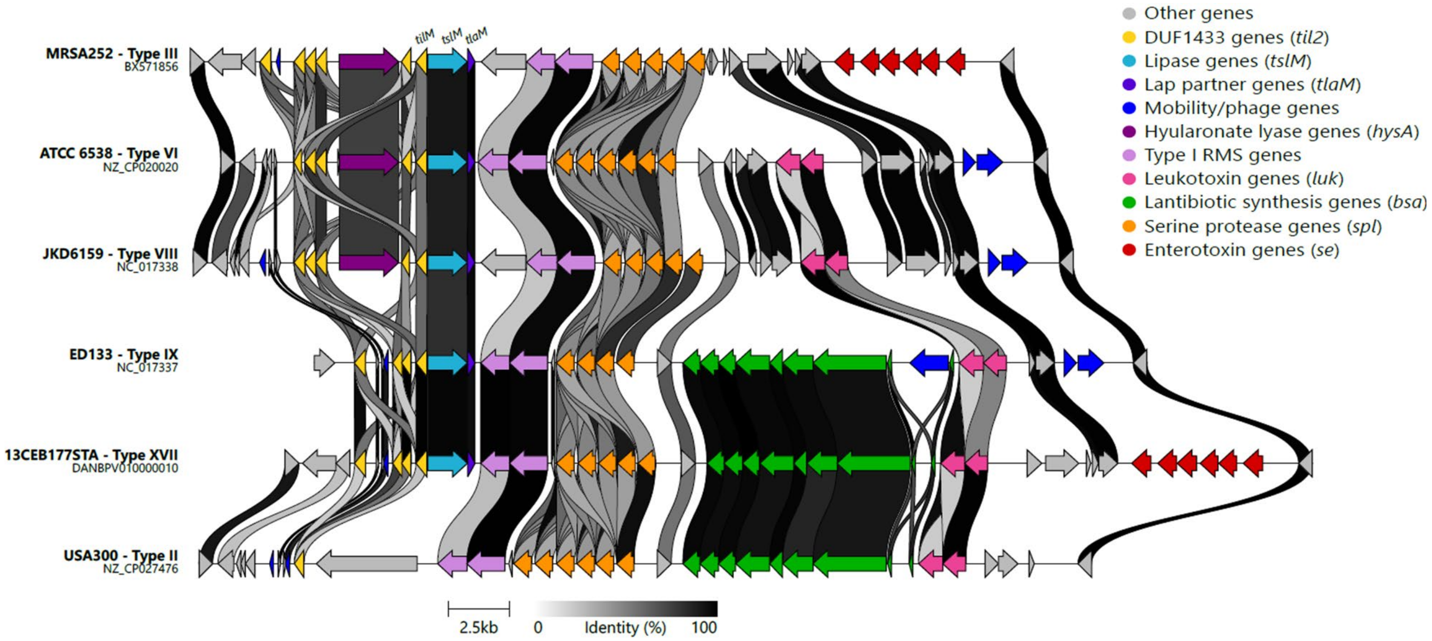


**Figure S5.** Alignment of vSaβ island subtypes that encode *tslM*. vSaβ genomic island subtypes III, VI, VIII, IX and XVII showing *tslM* (light blue) alongside *tilM* and *tlaM*. Also shown is the type II island from USA300, which does not encode *tslM* but still encodes an "orphan" Til2/DUF1433 (yellow). The majority of other vSaβ subtypes encode at least one Til2 (not shown but see (Kläui et al., 2019)). Alignment generated using Clinker.

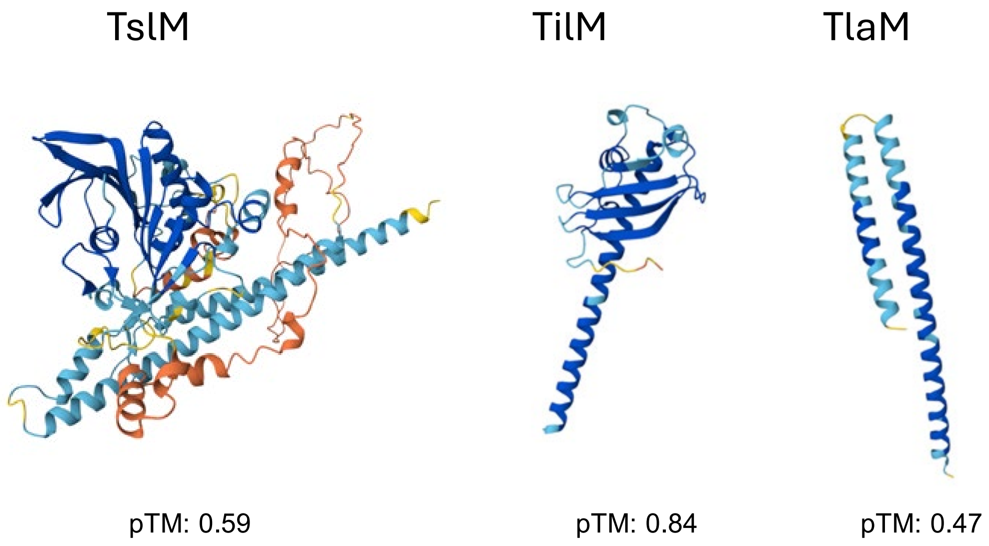


**Figure S6.** Alphafold3 models of TslM, TilM and TlaM coloured by confidence (where red is low confidence and blue high confidence). Predicted Template Modeling scores (pTM) are also given.

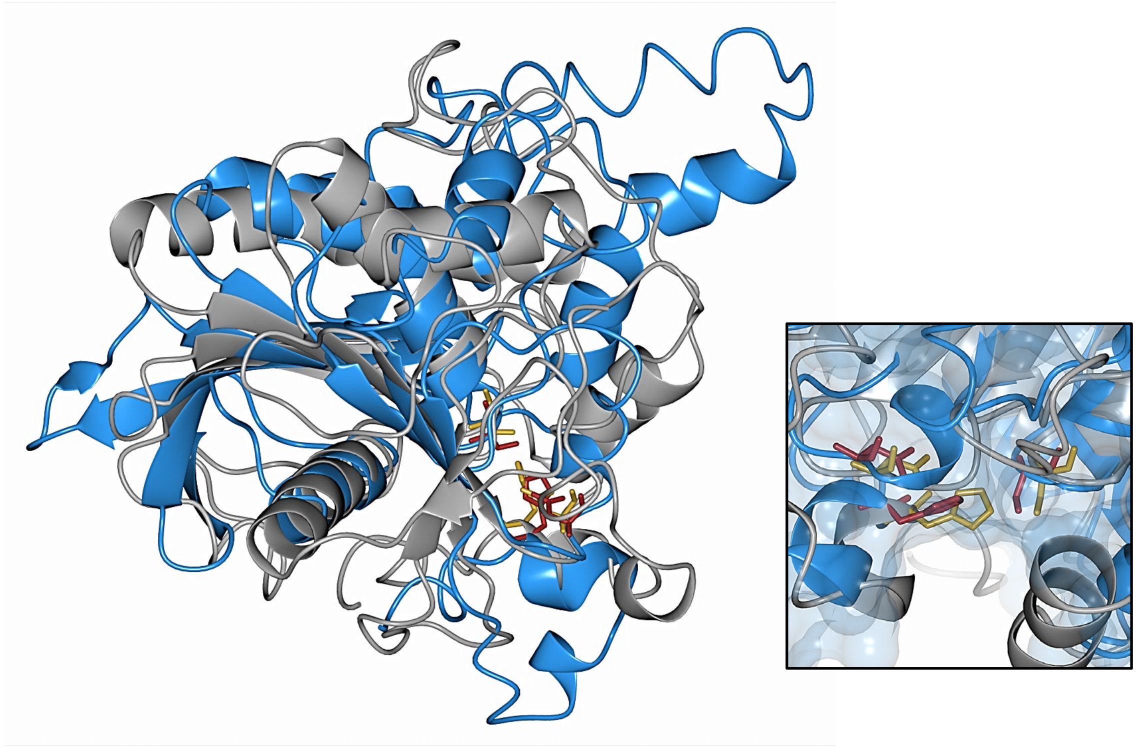

**Figure S7.** Lip2 and TslM superposition. Superposition of the lipase domain of TslM (residues 1-268, blue) and Lip2, a characterised lipase (PDB ID: 3o0d(Bordes et al., 2010), grey). The predicted catalytic triad of TslM (S171A, D226A, H261A) is shown in red, and the catalytic triad of Lip2 (S162, D230, H289) in gold. Inset: close up view of the predicted active site of TslM superposed with that of Lip2, with surface view shown. Superposition was performed with the SSM method.

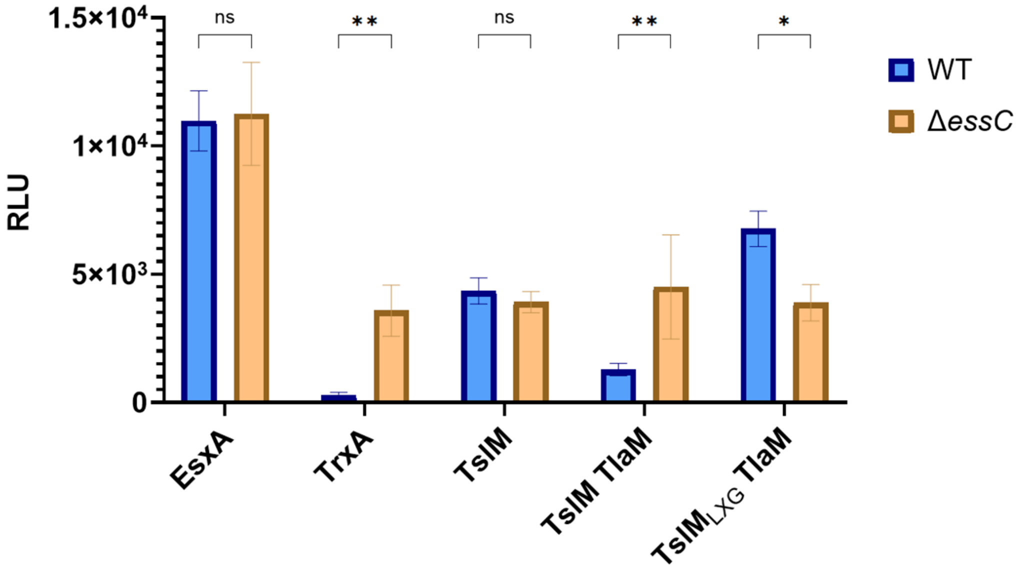


**Figure S8.** Nanoluciferase assay of cytoplasmic fraction of samples from Figure 4. Relative luminescence of cytoplasmic fraction from cultures producing C-terminal pep86-tagged TslM alone, or with TlaM, or a pep86-tagged C-terminal domain alongside TlaM (TslM$_{LXG}$-TlaM) in both wild type (blue) and Δ*essC* (orange) strains. C-terminally pep86-tagged EsxA (T7SS-secreted substrate) and TrxA (cytoplasmic protein) were included as controls. Readings were taken at maximum luminescence which reached 7 minutes following the addition of luciferase substrate and 11S. Error bars show standard deviation of three biological replicates. Significant difference was determined by 2way ANOVA: P values represented by asterisks: * <0.05, ** <0.01.

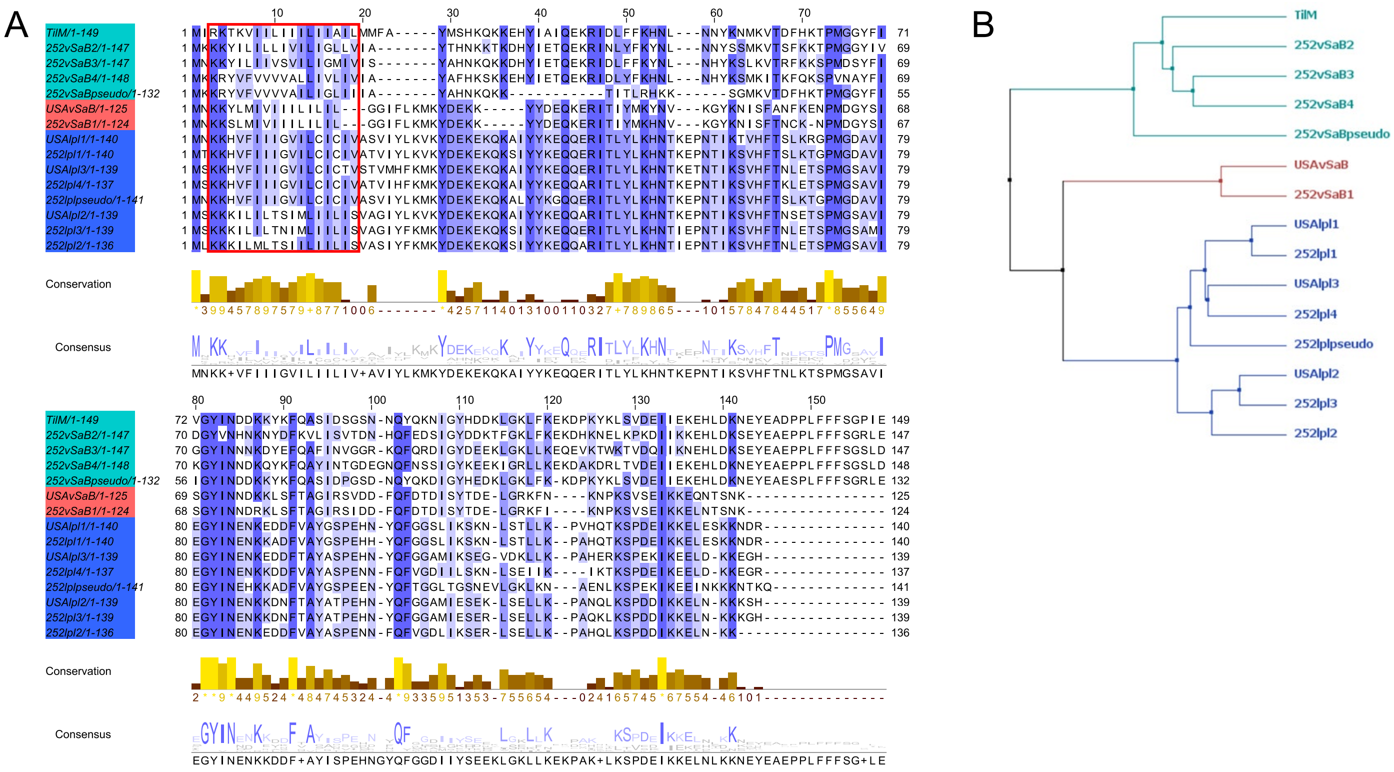


**Figure S9.** Multiple sequence alignment of the Til2 proteins encoded by USA300 and MRSA252. A. Alignment performed with the Muscle algorithm, visualised in Jalview. Residues and logo coloured by sequence conservation. *til2* genes are numbered from the first gene in a downstream direction in their respective loci: vSaβ or *lpIIII*. Pseudogenes (indicated) were repaired for the purposes of the alignment by the manual removal of frameshifts and subsequent translation to the shown amino acid sequence. The conserved N-terminal transmembrane helix is boxed in red. Sequences are coloured and grouped according to the tree shown in B. B. Maximum likelihood phylogeny generated from the 15 Til2 proteins encoded in USA300 and MRSA252. Three clear clades emerge, with TilM grouping best with the other Til2 proteins encoded at the MRSA252 vSaβ locus, and all *lpIIII* encoded Til2 proteins from both strains grouping together. This indicates that immunity protein diversity is locus specific, possibly reflecting the specific evolution of immunities required to protect against toxin subtypes within each toxin family. NRS_RS09835, used as the noncognate Til2 in toxicity experiments of Fig 7B, is labelled here as USAvSaβ.

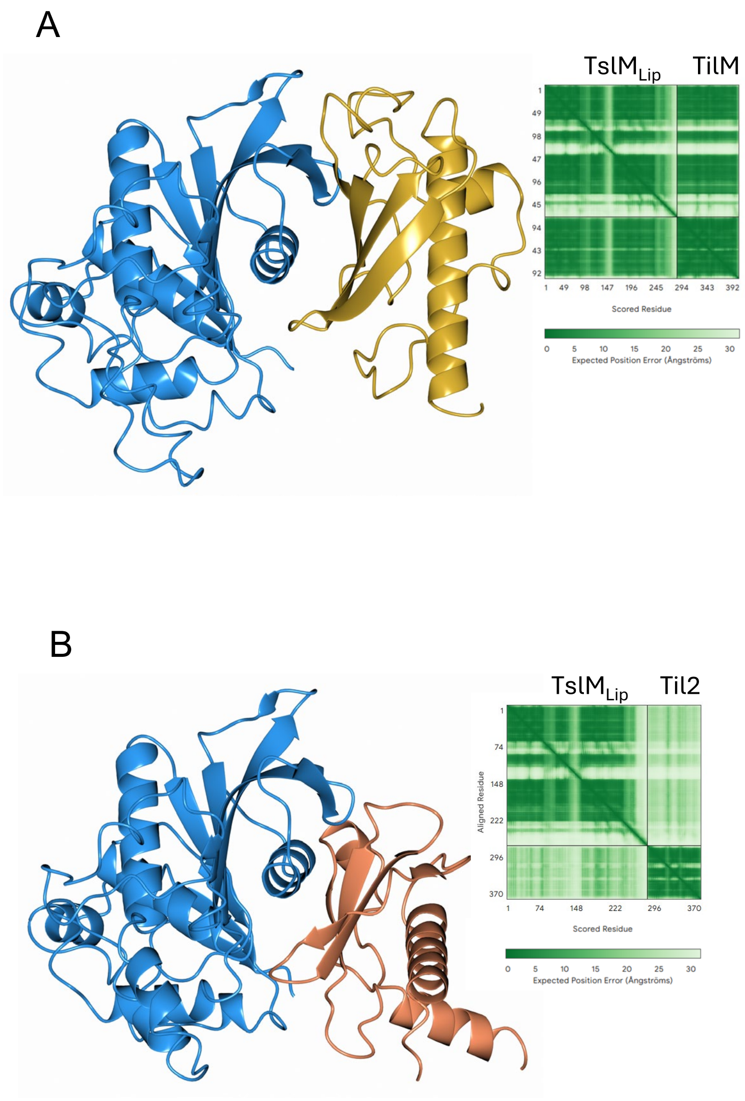


**Figure S10.** Modelling of Til2 binding to TslM. A. TslM lipase domain (blue) and TilM (without TM helix, gold) complex generated by AlphaFold3. Inset: Predicted Aligned Error plot of the complex generated by AlphaFold3, iPTM = 0.89. B. TslM lipase domain (blue) and noncognate Til2 NRS_RS09835 (without TM helix, orange) complex generated by AlphaFold3. Inset: Predicted Aligned Error plot of the complex generated by AlphaFold3, iPTM = 0.23. Note the difference in predicted binding position and ipTM confidence scores between the complexes.
